# ORFhunteR: an accurate approach for the automatic identification and annotation of open reading frames in human mRNA molecules

**DOI:** 10.1101/2021.02.05.429963

**Authors:** Vasily V. Grinev, Mikalai M. Yatskou, Victor V. Skakun, Maryna K. Chepeleva, Petr V. Nazarov

## Abstract

**Motivation:** Modern methods of whole transcriptome sequencing accurately recover nucleotide sequences of RNA molecules present in cells and allow for determining their quantitative abundances. The coding potential of such molecules can be estimated using open reading frames (ORF) finding algorithms, implemented in a number of software packages. However, these algorithms show somewhat limited accuracy, are intended for single-molecule analysis and do not allow selecting proper ORFs in the case of long mRNAs containing multiple ORF candidates.

**Results:** We developed a computational approach, corresponding machine learning model and a package, dedicated to automatic identification of the ORFs in large sets of human mRNA molecules. It is based on vectorization of nucleotide sequences into features, followed by classification using a random forest. The predictive model was validated on sets of human mRNA molecules from the NCBI RefSeq and Ensembl databases and demonstrated almost 95% accuracy in detecting true ORFs. The developed methods and pre-trained classification model were implemented in a powerful ORFhunteR computational tool that performs an automatic identification of true ORFs among large set of human mRNA molecules.

**Availability and implementation:** The developed open-source R package ORFhunteR is available for the community at GitHub repository (https://github.com/rfctbio-bsu/ORFhunteR), from Bioconductor (https://bioconductor.org/packages/devel/bioc/html/ORFhunteR.html) and as a web application (http://orfhunter.bsu.by).

## Introduction

High-throughput technologies allow capturing sequence information about whole transcriptomes in a reasonable time and cost. A number of high-performance computational approaches has been developed to restore the structure of full-length RNA molecules (or transcripts) from short RNA-Seq reads and to get qualitative and quantitative characteristics of these molecules (Mardis, 2017; Reuter, et al., 2015). One of the most important properties of transcripts is their coding potential that can be estimated using numerous algorithms. Methods implemented in NCBI ORFfinder (Sayers, et al., 2019), CPC2 (Kang, et al., 2017), OrfPredictor (Min, et al., 2005), ORFik (Tjeldnes and Labun, 2019), getORF (Rice, et al., 2000) are based on the selection of the longest open reading frame (ORF) from all possible candidates. Other tools, often used for metagenomics analysis, are based on vectorization of sequence features of ORF-candidates, which is an efficient conversion of nucleotide sequence into a vector of sequence features (Al-Ajlan and El Allali, 2018; Al-Ajlan and El Allali, 2019; El Allali and Rose, 2013; Hoff, et al., 2008; Trimble, et al., 2012; Zhang, et al., 2017). These features can be evaluated by a machine learning models in order to select true ORFs. Alternatively, statistical methods can be used to identify the most probable ORFs (Rainey and Repka, 2013). There is a spectrum of diverse algorithms implemented in the existing tools, such as ORF Investigator (Dhar and Kumar, 2018), GetOrf (Artimo, et al., 2012), OrfM (Woodcroft, et al., 2016), TransDecoder (Haas, 2018), findorf (Krasileva, et al., 2013), ORF Finder (Stothard, 2000), OrfPredictor Server (Min, et al., 2005), StarORF (Shubert, et al.), EasyGene (Nielsen and Krogh, 2005), FramePlot (Bibb, et al., 1984) or Third Position GC Skew Display (Ishikawa and Hotta, 1999). However, these tools have several limitations. First, they do not allow making a reasonable choice of one of the ORFs, if multiple candidates exist in the considered RNA molecule. Second, they usually work with individual molecules, and do not provide automated computational tools for a high-throughput analysis of large data sets. Finally, they show low accuracy predicting ORFs, require significant computing resources, considerably long computation time, and lack of integration with software aimed at the analysis of structural and functional characteristics of RNA molecules.

A promising way that could wave the main drawbacks of existing algorithms, is to select the informative parameters describing candidate ORF fragments using advanced algorithms of nucleotide sequence vectorization (Bao, et al., 2014; Mao, et al., 2014), and then to apply an optimal prediction algorithm and identify the most probable or true ORF sequence among candidates. This approach needs only a limited set of input features (here we used 104), which is significantly less than 4000-5000 considered earlier (Al-Ajlan and El Allali, 2018). The use of a random forest classifier (Breiman, 2001) has several advantages over more complex deep learning techniques (Al-Ajlan and El Allali, 2019; Wen, et al., 2019). A classical random forest classifier requires less computation power and a smaller training set, is less prone to an overfitting problem, and is much more interpretable.

Here we present a computational approach and its implementation, an ORFhunteR – R/Bioconductor package, aimed at automatic determination of true ORFs in mRNA molecules. The proposed method is based on i) vectorization of nucleotide sequences into sequence-dependent features and ii) a random forest classifier.

## Methods

### Data

We downloaded sequences of well-annotated 128161 mRNA molecules of protein-coding genes and 4235 long non-coding RNA (lncRNA) molecules from the manually curated NCBI RefSeq database (NCBI RefSeq release 109 based on GRCh38.p12 reference assembly of human genome). Coordinates and extracted sequences of highly confident true ORFs in mRNA molecules were collected, resulting in 113085 records in total. Additionally, we calculated coordinates and extracted 108800 sequences of pseudo-ORFs from lncRNA molecules. Similar to real ORFs, pseudo-ORFs begin with ATG start codon and end in-frame with one of the stop codons, but are not translated into proteins. These two sets of ORFs were combined into a single well-balanced discovery data set of true ORFs and pseudo-ORFs (imbalance index of 1.04) (Orriols-Puig and Bernadó-Mansilla, 2008).

In addition, we downloaded Ensembl annotations of human genes (Ensembl release 97 based on GRCh38.p12 reference assembly of human genome) and extracted the sequences of mRNA and lncRNA molecules. To avoid artifacts of Ensembl annotation algorithm, we excluded: i) mitochondrial transcripts, ii) 5’ incomplete transcripts, containing canonical stop codon but lacking a start codon inside the sequence, iii) 3’ incomplete transcripts containing canonical start codon ATG but lacking a stop codon inside the sequence, iv) both 5’ and 3’ incomplete transcripts lacking start and stop codons inside the sequence, v) and transcripts with non-canonical start codons CTG, GTG or TTG. We combined filtered Ensembl mRNAs (56765 records in total) and lncRNAs (74980 records in total) into a single test data set of RNA molecules.

### Overview of the computational approach

The proposed computational approach for the automatic identification of the true ORFs integrates algorithms for vectorization (Zakirava, et al., 2019) and random forest-based classification (Breiman, 2001). Our pipeline is presented in Figure 1, and includes the following five steps: building a discovery data set of ORFs, vectorization of discovery data set ORFs into sequence features, training of the classification model, identification of the true ORFs in a set of mRNA molecules, and (optionally) annotation of the identified ORFs.

**Figure 1.**
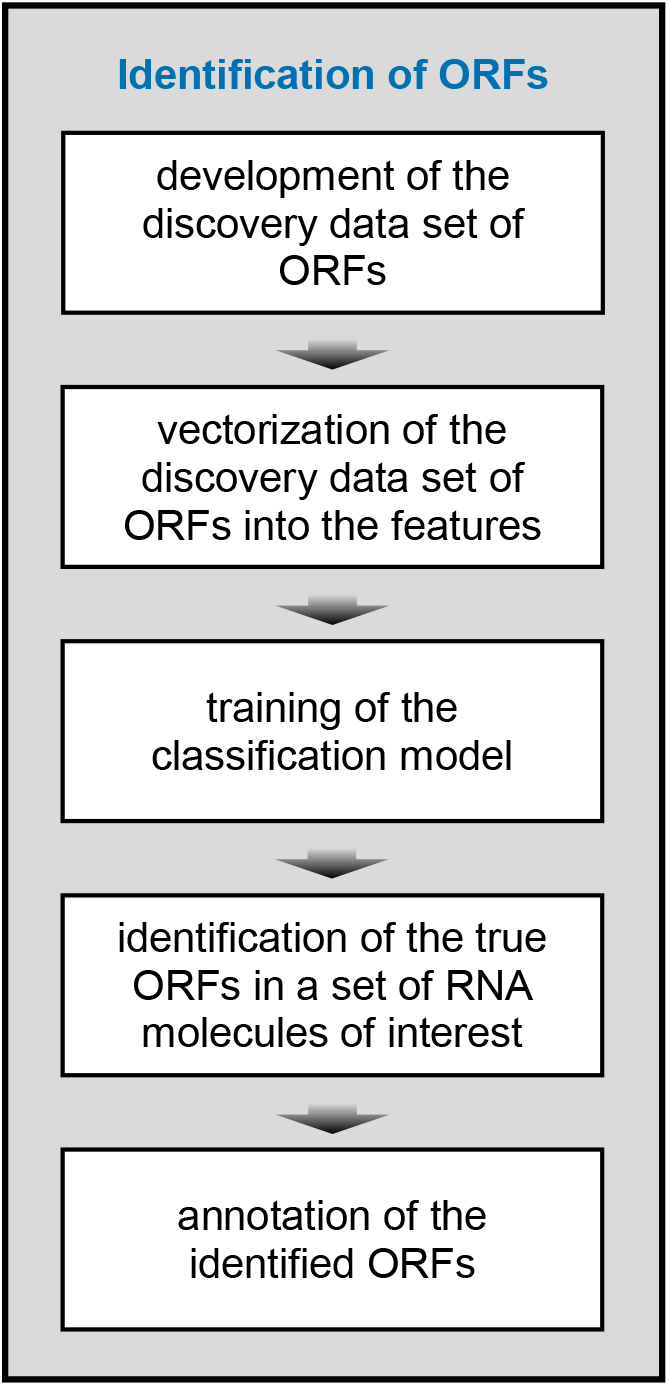
The main stages for automatic identification of ORFs in RNA molecules.

The discovery data set of ORFs was developed as described in previous section. These ORFs were vectorized into 104 sequence features: frequency of mono-, di- and trinucleotides (84 features) (Zakirava, et al., 2019), nucleotide correlation factors (Mao, et al., 2014), sequence lengths (8 features in total), and parameters of the Category-Position-Frequency (CPF) model representing the local frequency-based entropy values of sequences (12 features) (Bao, et al., 2014). As was shown in numerous studies, these features are indeed informative for classification of coding and non-coding regions of RNA molecules (Bao, et al., 2014; Mao, et al., 2014; Zakirava, et al., 2019). Below the 104 features are considered in details.

The frequency parameters of mono-, bi- and trinucleotides were calculated by the standard algorithms, integrated in the R/Bioconductor package Biostrings.

When calculating the parameters of the nucleotide correlation factors, the sequence is divided into *m* sections with a length of 20 base pairs. The probabilities 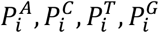, or abovementioned frequencies, of four mononucleotides A, C, T, G in each *i*-th section are calculated. Six correlation factors of nucleotide pairs are:

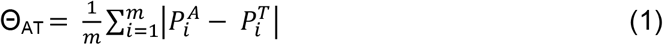

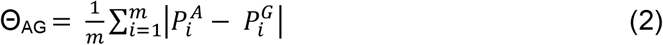

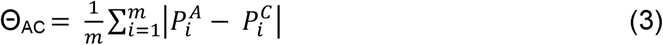

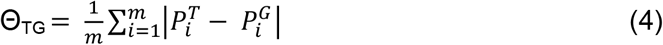

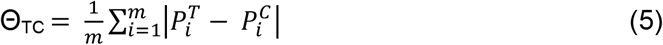

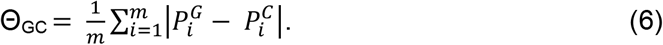

The sequence length-dependent features are represented by both the number of nucleotide between the start and the stop codons and its logarithm.

In the CPF model (Bao, et al., 2014), the nucleotide sequence is served as a parent of three sequences of the same length, containing two possible symbols instead of four bases. The transformation is carried out according to the chemical structure of nucleotides: the substitution in the first sequence is based on the fact that the nucleotides A and G contain a purine group, and the bases T and C – a pyrimidine group. The second sequence takes into account that the nucleotides A and C involve an amino group and T and G – a ketamine group. When constructing the third sequence, the features of complementary bonds are taken into account: the A-T bond is weak, and the G-C bond is strong. Further, from each sequence, 4 new binary sequences are generated (Bao, et al., 2014). The binary sequences are translated into the sequences of the local frequencies *LF^w^_r_*:

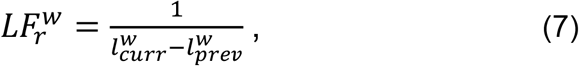

where *r* is the order of occurrence of a word *w*, *r* = 1, 2, .., *n, n* is the binary sequence length, *l^w^_r_* denotes the position of the *r*-th occurrence of the *w*, and *l^w^_0_* is defined as 0.

The sequence of local frequencies is converted into a sequence of a partial sum:

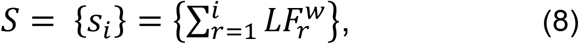

where *i* = 1, 2, .., *n*. The entropy is calculated as:

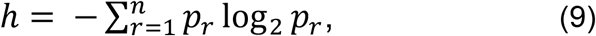

where

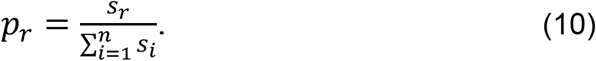

The resulting entropy is the feature value of the binary sequence. Entropy is calculated for each of 12 binary sequences, representing 12 features generated by the CPF model.

With above developed and vectorized discovery data set of ORFs, the 104 parameters for a random forest classifier were calculated. The random forest classification algorithm was selected since it has a high classification accuracy, quite stable against overfitting, provides estimation of a feature informativity based on a selected criterion, it is fast and requires less computational resources comparing to advanced deep learning algorithms (Wen, et al., 2019).

Next, the trained classification model was applied for identification of the true ORFs among ORF candidates extracted from mRNA molecules of interest. For each mRNA molecule, ORF candidates were inferred as sequences beginning from ATG start codon and ending in-frame with one of the stop codons. These ORF candidates were vectorized into 104 sequence features and used as input in classification model for identification of the true ones. Finally, identified ORFs could be annotated using various metrics implemented in our software.

### Computations and program package realization

We implemented computational algorithms in the R and C++ programming languages using R/Bioconductor and CRAN packages. Usage of C++ with Rpp package increased the performance of the analysis by almost 100-fold.

To develop the classifier, based on the random forest method, the R package randomForest was used (Breiman, 2001). One of the advantages of this implementation is its ability to assess the significance of the features by the Gini index (Hastie, et al., 2009). We tested random forests composed of 200, 300, 400, 500, and 5000 trees. The classification error for these trees varied less than by 0.5%. The computation time increased almost linearly with an increase of the number of trees. This also led to increase of the file size, obtained when saving the model. A decision was made by voting among 500 trees, which provides high accuracy with reasonable computation time.

All computations and data analyzes were carried out on a PC with a 12-core Intel i9 processor (3.9 GHz), 64 Gb DDR4 RAM. The most time consuming step was training of the classification model, which took about 5 hours.

The developed algorithms are integrated into an ORFhunteR package (identification of the <u>O</u>pen <u>R</u>eading <u>F</u>rames by <u>hunt</u>ing with <u>R</u> codes). Several versions of ORFhunteR software packages were implemented, including the basic R-package deposited in GitHub repository (https://github.com/rfctbio-bsu/ORFhunteR) and online application (http://orfhunter.bsu.by).

The functional structure of the ORFhuneR package is shown in Figure 2. From this package, the function *loadTrExper* loads a set of input transcripts. In the uploaded transcripts, ORF candidates are identified using the functions *codonStartStop* and *findORFs*. These ORF candidates are vectorized into sequence features using the R function *vectorizeORFs* in conjunction with the C++ functions *getBaoMetrics* and *getCorrelationFactors*. The vectorized ORF candidates are classified into true ORFs and pseudo-ORFs by function *predictORFs*. If necessary, the nucleotide sequence of the true ORFs can be obtained using the function *getSeqORFs*. Finally, identified true ORFs can be annotated by function *annotateORFs* in conjunction with the functions *findPTCs* and *translateORFs*. Herewith, the function *findPTCs* identifies premature termination codons in transcripts of interest while function *translateORFs* translates ORFs to proteins.

**Figure 2.**
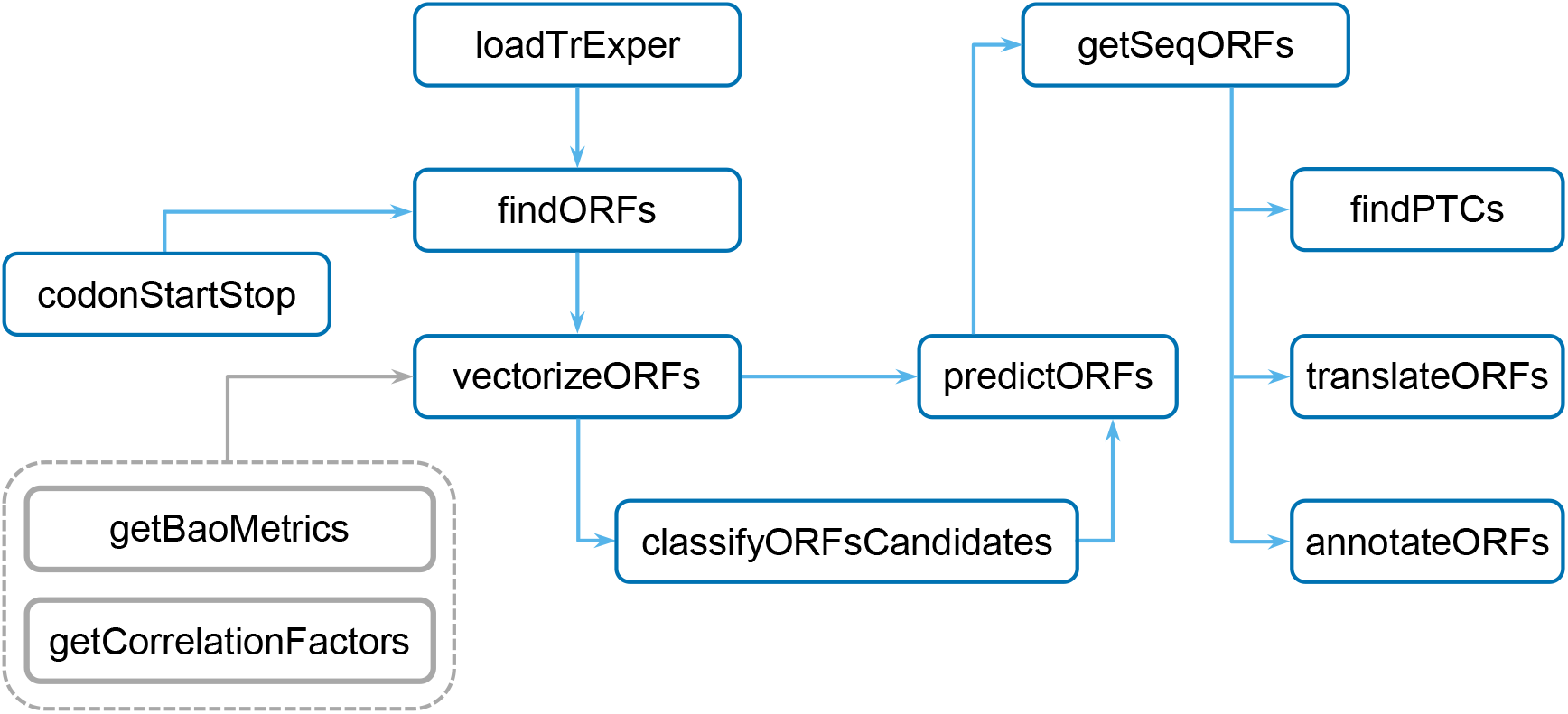
The functional structure of ORFhunteR package.

## Results and discussion

For training of the classification model, we utilized sequence data from NCBI RefSeq database due to a number of minimal technical artifacts comparing to alternative sources of sequence information. Collected sequences were used for development of the discovery data set of ORFs. We next vectorized these ORFs into 104 sequence features and randomly divided them into training (75% of records) and validation (25% of records) sub-sets of sequences. Internal evaluation of the classification accuracy was performed on the validation sub-set.

To assess the classification accuracy of true ORFs and pseudo-ORFs, the accuracy was calculated as the percentage of correctly classified ORFs to the total number of inputted ORFs. In our classification model, the classification accuracy of the validation sub-set of ORFs reached 99.4%. Nearly the same value was obtained for the entire discovery data set. These results were in a very good agreement with principal component analysis, according to which the two classes of ORFs are well separated (Figure 3a). Herewith the parameters of the CPF model and ORF length turned out the most important sequence features in the class determination of ORFs (Figure 3b). The less informative sequence features are the nucleotide frequencies and nucleotide correlation factors.

**Figure 3.**
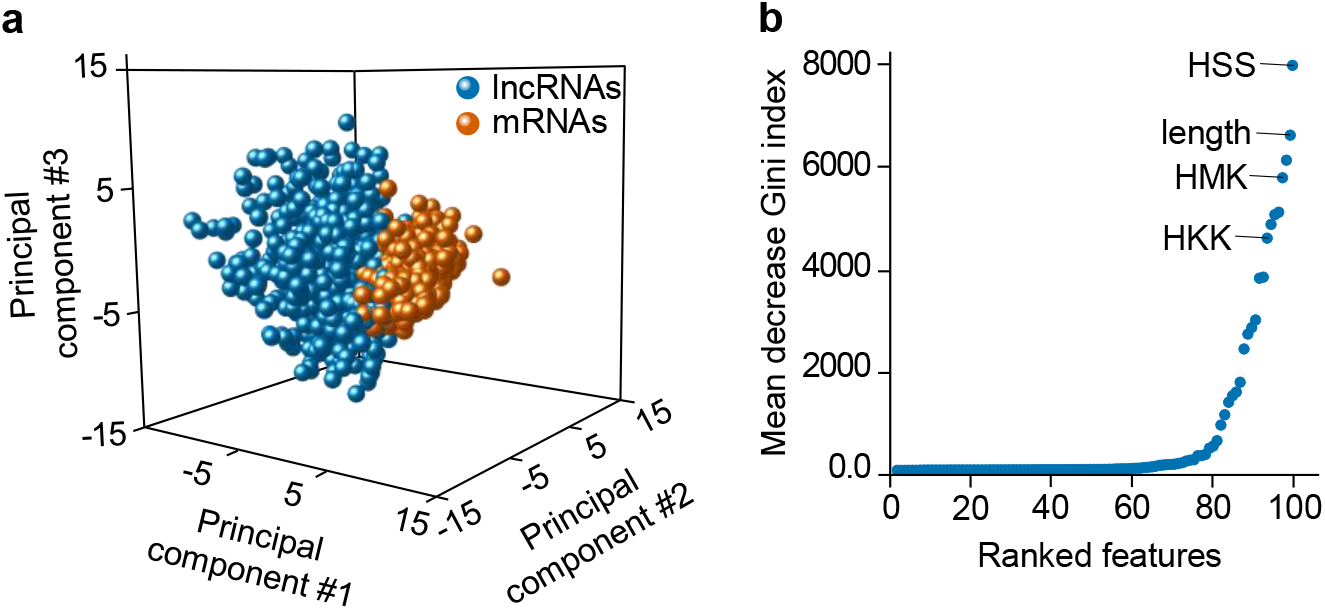
The sequences features of the discovery data set of ORFs. **(a)** LncRNAs and mRNAs derived references ORFs in the space of the first three principal components.For each class of RNA molecules, only 1000 ORFs were randomly selected for compact representation. **(b)** The importance of sequence features in determining the class of ORF. The importance of features was assessed with Gini index in a random forest-based classification procedure. The higher mean decrease in the Gini index characterizes the quality of the tree node partitioning (the lower the value, the better the node partitioning), the more informative the feature is. The HSS, HMK and HKK features are the examples of CPF model parameters (Bao, et al., 2014).

To validate our classification model further, we calculated coordinates and extracted the sequences of all possible variants of ORF candidates from NCBI RefSeq and Ensembl human mRNA molecules as described above.

Our classification model demonstrated 98.3% accuracy in identification of true ORFs in NCBI RefSeq data. Understanding that the classification model was constructed based on the true ORFs and pseudo-ORFs generated from the NCBI RefSeq RNAs that might lead to an overfitted model, we then analyzed the Ensembl RNA data. On this dataset, the approach allows to identify the true ORFs with an accuracy of 94.9%. In fact, 91.9% of ORFs that were identified in Ensembl human mRNA molecules demonstrates probability ≥ 0.9 for the “winning” class of true ORFs (Figure 4a, b; the probability of the ORF to be coding is calculated by the random forest classifier). At the same time, distribution of probability values for pseudo-ORFs from long non-coding RNA molecules differs (Figure 4c).

**Figure 4.**
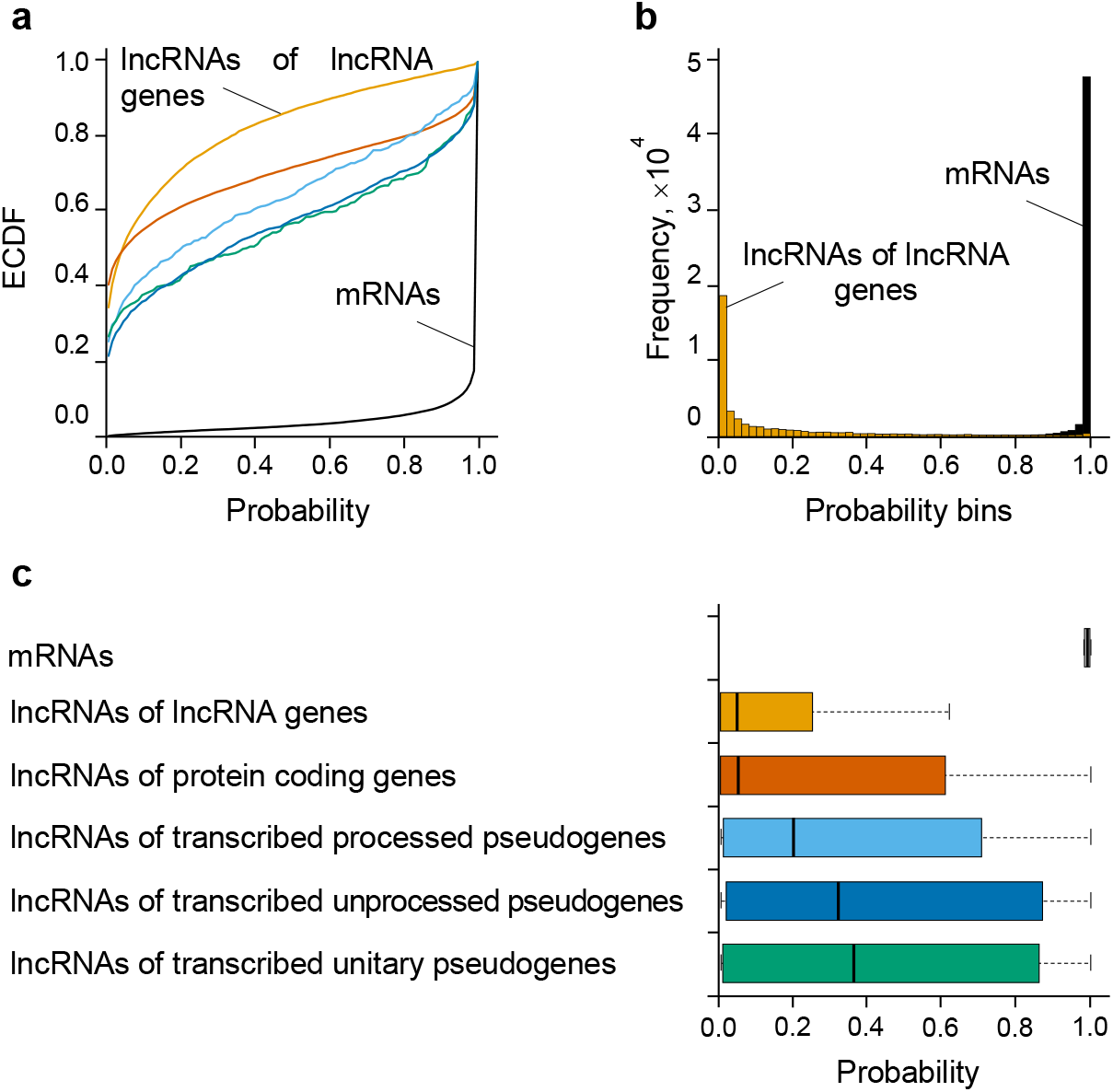
Distribution of probability values on identification of true ORFs and pseudo-ORFs in different human RNA molecules (Ensembl release 97, GRCh38.p12 human reference genome assembly). **(a)** Empirical cumulative distribution of probability values for ORFs that were identified in mRNAs (**-**) and lncRNAs encoded by lncRNA genes 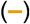, protein-coding genes 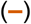, transcribed processed pseudogenes 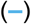, transcribed unprocessed pseudogenes 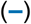 and transcribed unitary pseudogenes 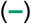. **(b)** Frequency of probability values for ORFs that were identified in mRNAs and lncRNAs encoded by lncRNA genes. **(c)** Boxplot demonstrating the distribution of probability values for ORFs that were identified in mRNAs and lncRNAs different gene biotypes.

However, choosing of a suitable threshold of probability to discriminate true ORFs and pseudo-ORFs in mRNA molecules is still challenging. We suggest two approaches for solving this problem. The first approach is to choose an adaptive probability threshold at which the rate of detection of true ORFs is maximized and the rate of detection of pseudo-ORFs is minimized. With the Ensembl dataset, this threshold value can be 0.9 (Figure 5). Alternative approach combines classifying the coding RNA molecules at first and then determining the ORFs for those coding RNAs at second, meaning no needs for using a probability threshold. The first step of this combined approach can be done in independently using, for example, convolutional neural networks (Wen, et al., 2019).

**Figure 5.**
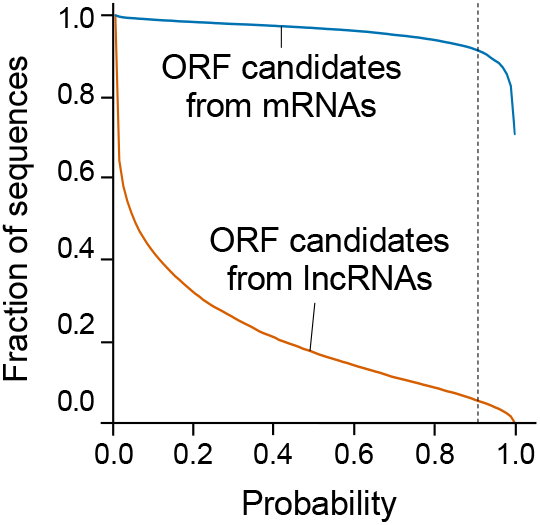
Relationship between fraction size of ORFs and probability of true ORF class for Ensembl-based human mRNAs and lncRNAs.

## Conclusion

The efficient computational approach for the identification of unknown ORFs in mRNA molecules was developed and integrated into the corresponding R/Bioconductor package ORFhunteR. It is based on vectorization of the sequence features of ORFs candidates and predicting the most evident by a random forest classifier. Our numerical tests resulted the accuracy of ORFs identification 98.3% and 94.9% on the discovery and test data sets. In our mind, the developed approach has three advantages over the competing strategies based on neural networks (Al-Ajlan and El Allali, 2019; Wen, et al., 2019): i) it requires less computing resources and works out much faster; ii) it is less prone to overfitting as a neural network and uses a limited set of vectorized features (unlike the statistical approaches using thousands of features); iii) finally, random forest classifiers show way better interpretability compared to deep learning or boosting models.

## Code availability

Code developed in this study is freely available via GitHub project ORFhunteR at https://github.com/rfctbio-bsu/ORFhunteR.

## Acknowledgements

The work of V.V.G., M.M.Y., V.V.S. and M.K.S. was supported by the Ministry of Education of the Republic of Belarus, grant GPNI “Convergenciya–2020” N3.08.3 (registration number 20190531). P.V.N. was supported by Luxembourg National Research Fund (C17/BM/11664971/DEMICS).

